# Relationships between pond water and tilapia skin microbiomes in aquaculture ponds in Malawi

**DOI:** 10.1101/2021.12.06.470702

**Authors:** Jamie McMurtrie, Shayma Alathari, Dominique L. Chaput, David Bass, Camerson Ghambi, Joseph Nagoli, Jérôme Delamare-Deboutteville, Chadag Vishnumurthy Mohan, Joanne Cable, Ben Temperton, Charles R. Tyler

## Abstract

Intensification of fish farming practices is being driven by the demand for increased food production to support a rapidly growing global human population, particularly in lower-middle income countries. Intensification of production, however, increases the risk of disease outbreaks and thus the likelihood for crop losses. The microbial communities that colonise the skin mucosal surface of fish are poorly understood, but are important in maintaining fish health and resistance against disease. This skin microbial community is susceptible to disruption through stressors associated with transport, handling and the environment of intensive practices, and this risks the propagation of disease-causing pathogens. In this study, we characterised the microbial assemblages found on tilapia skin — the most widely farmed finfish globally — and in the surrounding water of seven earthen aquaculture ponds from two pond systems in distinct geographic regions in Malawi. Metabarcoding approaches were used to sequence the prokaryotic and microeukaryotic communities. We found 92% of prokaryotic amplicon sequence variants were common to both skin and water samples. Differentially enriched and core taxa, however, differed between the skin and water samples. In tilapia skin, *Cetobacterium, Paucibacter, Pseudomonas* and Comamonadaceae were enriched, whereas, the cyanobacteria *Cyanobium, Microcystis* and/or *Synechocystis*, and the diatom *Cyclotella*, were most prevalent in pond water. Ponds that clustered together according to their water prokaryotic communities also had similar microeukaryotic communities indicating strong environmental influences on prokaryotic and microeukaryotic community structures. While strong site-specific clustering was observed in pond water, the grouping of tilapia skin prokaryotes by pond site was less distinct, suggesting fish microbiota have a greater buffering capacity against environmental influences. The characterised diversity, structure and variance of microbial communities associated with tilapia culture in Malawi provide the baseline for studies on how future intensification practices may lead to microbial dysbiosis and disease onset.

**Highlights:**

- Fish skin and pond water communities differ structurally, but share common taxa
- Pond locations have a stronger influence on water *versus* fish skin microbiome community structure
- Selected skin-associated taxa could be used to monitor dysbiotic events in aquaculture
- Taxa with opportunistic pathogen potential were identified at low abundance

## 1. Introduction

Capture fisheries will not be able to satisfy the demand for seafood products from an ever-increasing human population with rising living standards (Henchion et al., 2017) combined with plateauing, and in some cases declining, wild fish stocks due to overfishing and ecosystem degradation (Link and Watson, 2019). Seeking to meet this demand for aquatic products, many aquaculture farming practices are undergoing intensification. Shifting from extensive to intensive and semi-intensive practices in aquaculture, however, is often associated with increased incidence of infectious disease (Hinchliffe et al., 2020; Pulkkinen et al., 2010). Intensification can cause chronic stress that adversely impacts fish physiology resulting in reduced growth and impaired disease resilience. Increasing pond stocking rates and levels often occurs with insufficient amounts of clean water, leading to the deterioration of water quality, including dissolved oxygen, pH and ammonia (Abdel-Tawwab et al., 2014; Sundh et al., 2019), which in turn impacts negatively on fish growth and health, and renders the fish more susceptible to diseases. Regular restocking of ponds with fish of uncertain health status to compensate for mortalities, in turn, increases the likelihood of repeated introductions of sub-clinical infections (Bondad-Reantaso et al., 2005; Murray and Peeler, 2005).

Disease remains a huge challenge for aquaculture, particularly in Asia where 89% of global aquaculture production occurs (FAO, 2020c). Successful management of disease risk and intensification of aquatic species production requires a better understanding of the relationships between the microbial systems (microbiomes) of both the cultured aquaculture species and of the environments in which they are grown (Bass et al., 2019). The study of microbiomes in aquaculture is gaining momentum and recent studies have investigated how pond and fish treatments (e.g. antibiotics, dietary supplements, probiotic treatments and pond fertilisers) affect fish microbiomes (Limbu et al., 2018; Minich et al., 2018; Suphoronski et al., 2019; Tan et al., 2019). Much of this research has focused on the gut microbiome due to its intricate role in gut health, which when optimised can maximise feed conversion, growth, and overall aquaculture productivity (Perry et al., 2020). When considering disease resistance and/or susceptibility in fish aquaculture, however, arguably the microbial communities harboured on/in the skin and gills are likely to be equally if not more important.

These outer facing mucosal surfaces are in continuous contact with the aquatic environment and provide a primary barrier against invading pathogens (Legrand et al., 2018; Rosado et al., 2019b). The microbes colonising this skin niche include those specifically adapted to the host mucosal surface, as evidenced by host-species specificity of microbiome composition (Doane et al., 2020), but also microbes derived from the surrounding water community (Krotman et al., 2020). Relatively little is known about the environmental and host contributions to these microbial assemblages, particularly in aquaculture ponds. It is known, however, that skin colonisers have a direct connection with the host immune system helping to shape its function and responses (Kanther et al., 2014). Equally, the immune system provides feedback in sculpting the microbial community structure (Kelly and Salinas, 2017; Tarnecki et al., 2019). If these finely balanced communities are disrupted, to a state known as dysbiosis, resulting health complications and disease may occur. The fish skin microbiome has been reported to change following stressful events, such as high stocking densities and hypoxia (Boutin et al., 2013), in fish showing clinical signs of gastrointestinal enteritis (Legrand et al., 2018) and also following viral infection (by salmonid alphavirus; see Reid et al., 2017), bacterial infection (by *Photobacterium damselae*; see Rosado, Xavier, et al., 2019b) and macroparasitism (by the sea lice *Lepeophtheirus salmonis;* see Llewellyn et al., 2017). In all of these cases, there was a decrease in abundance of reputedly beneficial taxa, concurrent with an increase in opportunistic pathogens. The resulting theory is that dysbiosis within the skin microbiome causes fish to become more susceptible to secondary bacterial infections. This has been shown for exposure to the antimicrobials rifampicin in *Gambusia affinis* Baird & Girard (Carlson et al., 2015) and potassium permanganate (Mohammed and Arias, 2015) in *Ictalurus punctatus* Rafinesque, where increased mortality occurred for dysbiotic fish compared with controls when challenged with the disease-causing *Edwardsiella ictaluri* and *Flavobacterium columnare*, respectively.

A limitation in the majority of microbiome studies, regardless of host species, is a focus on the bacterial community only with little or no attention given to the remaining microbial community members. This includes microeukaryotes, a taxonomic group that encompasses protists, microfungi, microalgae, and microbial metazoans (Bass and del Campo, 2020; del Campo et al., 2019), as well as viruses that infect an expansive host range including microeukaryotes, bacteria and the animal host (Gadoin et al., 2021). Microeukaryotic communities are well described in some settings, such as the contribution of microalgae to primary production in the ocean (Benoiston et al., 2017). The relationships between microeukaryotes and animal hosts have predominantly focussed on parasitism and pathogenesis, yet microeukaryotes play an intricate role in the broader microbial community of host-associated niches. One of the best described examples is *Blastocystis*, a protist commonly found to colonise the gut of humans and other animal hosts. Its presence is thought to correlate with protection against several gastrointestinal inflammatory diseases by interacting with the bacterial community to promote a healthy microbiome (Laforest-Lapointe and Arrieta, 2018), specifically via an associated increase in bacterial diversity and strong co-occurrence patterns with reputed beneficial bacteria (Audebert et al., 2016; Beghini et al., 2017). The full role *Blastocystis* plays in human health remains unresolved and controversial. The extent of interactions occurring between bacteria and microeukaryotes and/or viruses in the fish skin microbiome is largely unknown and unreported.

Tilapia are the most widely farmed finfish in global aquaculture, produced in over 170 countries. Numerous species of tilapia are farmed, dominated by Nile tilapia (*Oreochromis niloticus* L.), and predominantly in lower-middle income countries (LMICs) across the Southeast Asian, African and South American continents (FAO, 2020b). Given their fast growth, adaptability to a variety of environmental culture conditions, and resilience against both disease and poor water quality, tilapia are now a production staple for many LMICs, and colloquially is often referred to as the aquatic chicken (FAO, 2020a). While some aquaculture species grown in LMICs, such as shrimp, are high-value products for export, the bulk of tilapia production is for domestic markets. As a consequence, fewer regulations exist for tilapia production (El-Sayed, 2019) and there has been far less scientific research for optimising sustainable production compared to some other high-value teleost species, such as Atlantic salmon.

Aquaculture in Malawi is in its relative infancy compared with other countries in Africa and Asia. Nevertheless, production has seen on average a 24% yearly growth between 2006 and 2016 (CASA, 2020). The levels of intensification or disease incidence seen in Malawi are low compared with Asia, but as demand increases, disease levels will inevitably increase also. Tilapia species cultured in Malawi include *Coptodon rendalli* Boulenger and *Oreochromis shiranus* Boulenger, with the notable absence of Nile tilapia, which is considered an invasive species. To fully elucidate the influence of microbiomes on fish health during disease processes, we need to better understand the relationships between the microbial diversity, community variance and structure in the mucosal surfaces of fish and those in the aquatic environment, including microeukaryotes (often excluded from microbiome studies), for disease-free populations. In this study therefore, we applied high throughput DNA sequencing for metabarcoding of the 16S and 18S ribosomal RNA (rRNA) small subunit (SSU) marker genes (which are conserved within prokaryotes and eukaryotes, respectively), to characterise the microbial communities of pond water and tilapia skin (*C. rendalli* and *O. shiranus*) from earthen aquaculture ponds in Malawi. With these data, we investigated the relationships between the pond water and skin microbiome. We identified differentially enriched and core taxa within the tilapia skin microbiome that are likely to play an important biological role for the host and may provide notable taxa for future studies to interpret disease events.

## 2. Materials and methods

### 2.1. Sample collection

Seven tilapia aquaculture earthen ponds were sampled in October 2017 from two pond systems in Malawi. Two ponds from a commercial farm were located in Maldeco, and a five ponds from a community pond syndicate were located 200 km further south in Blantyre (Supplementary Fig. 1). Two sample types were collected: pond water and tilapia skin swabs (Table 1). Pond surface water was collected from five locations within each pond by passing 200 mL of water through a polycarbonate filter (0.4 μm pore, 47 mm diameter, Whatman). The volumes of water filtered were affected by the amount of organic/particulate matter in the samples such that volumes were sampled until filters became saturated and prevented further filtration. Mucosal skin samples of tilapia flanks (*C. rendalli* and *O. shiranus*) were collected by swabbing three times along the entire length of the lateral line (Delamare-Deboutteville et al., 2021) with sterile polyester swabs (Texwipe). Filters were preserved in 1.8 ml of 100% molecular grade ethanol (FisherScientific), while swabs were preserved in 1.8 ml of RNA*later* (Qiagen), and stored at ambient temperature until transferred to the UK for prolonged storage at -20 °C, until used for DNA extraction and sequencing.

**Table 1.** Details of pond sites and samples for the pond water and fish skin swabs obtained from Malawian tilapia aquaculture ponds.

| Pond system location | Pond site | No. pond water samples | No. fish skin swabs * | Cultured species | Mean fish length (mm) ± SD | No. fish measured for length |
| --- | --- | --- | --- | --- | --- | --- |
| Blantyre, Malawi | 1 | 5 | 2 | *Coptodon rendalli* | 115 ± 11.8 | 5 |
| Blantyre, Malawi | 2 | 5 | 1 | *Coptodon rendalli* | 125 ± 19.1 | 5 |
| Blantyre, Malawi | 3 | 5 | 7 (5) | *Coptodon rendalli*,<br>*Clarias gariepinus* | 149 ± 9.3<br>N/A | 8 |
| Blantyre, Malawi | 4 | 5 | 8 | *Coptodon rendalli*,<br>*Oreochromis shiranus* | 137 ± 4.2<br>122 ± 6.6 | 5<br>3 |
| Blantyre, Malawi | 5 | 5 | 5 | *Coptodon rendalli* | 162 ± 18 | 5 |
| **Maldeco, Malawi** | 6 | 5 | 6 (4) | *Oreochromis shiranus* | 222 ± 30.5 | 5 |
| **Maldeco, Malawi** | 7 | 5 | 4 (3) | *Oreochromis shiranus* | 212 ± 12.7 | 5 |
| **Total** | **7** | **35** | **33 (28)** |  |  |  |

\*Numbers in parentheses refer to the number of 18S rRNA samples successfully sequenced, where this differs from 16S rRNA samples.

### 2.2. DNA extraction

Ethanol was removed from pond water filters by freeze-drying (ScanVac CoolSafe Pro, 4 L condenser at -110 °C) and filters were then stored at -80 °C. RNA*later* was removed from fish swab samples by vortexing the swabs for 30 seconds in 23 mL of 1x sterile phosphate buffered saline (Sigma) to allow detachment of microbes. The swab and solution were transferred to a syringe for filtration with a 0.22 μm Sterivex (Millipore) filter unit. Following ethanol and RNA*later* removal from filters and swabs, DNA was extracted with a CTAB/EDTA/chloroform method adapted from Bramwell et al. (1995) and Lever et al. (2015), and is available in full at (https://dx.doi.org/10.17504/protocols.io.bw8gphtw).

Briefly, for DNA extraction, filters were first suspended in 570 μl lysis buffer (30 mM Tris, 30 mM EDTA, pH 8, FisherScientific), freeze-thaw lysed in liquid nitrogen and homogenised by bead-beating with Lysing Matrix A Bulk Beads (Garnet) on the Qiagen TissueLyser II for 40 seconds at 30 Hz. The sample suspension was digested with 1 μL Ready-Lyse lysozyme (1000 U/ μL, Epicentre), and 3 μL proteinase K (20 mg/mL, Sigma) in 30 μL SDS (10% w/v, FisherScientific) for 1 hour at 55 °C. Samples were then incubated for 10 minutes at 65 °C in 120 μL NaCl (5 mM, Sigma) and CTAB solution (hexadecyltrimethylammonium bromide, 96 μL, 10% w/v, Sigma). An equal ratio of sample and 24:1 chloroform:isoamyl alcohol (Acros Organics) were used for extractions, with centrifugation at 14,000 x g, 4 °C for 5 minutes. The aqueous layer was retained for a second extraction, after which 1 μL of linear polyacrylamide solution (GenElute LPA, Sigma) was added to aid precipitation with 0.7 volumes isopropanol (Acros Organics). Following overnight incubation at 4 °C, samples were centrifuged at 21,000 x g, 4 °C for 30 minutes and the resulting pellet was washed with 70% ethanol (FisherScientific). After 10 minutes of centrifugation at 21,000 x g and pipetting off the ethanol, DNA pellets were resuspended in TE buffer (10 mM Tris-HCl, 1 mM EDTA, pH 8, Sigma) and stored at -20 °C until used for sequencing.

### 2.3. Metabarcoding

Metabarcoding of prokaryotic and microeukaryotic SSU rRNA marker genes was performed by PCR amplification with the Earth Microbiome Project recommended primers. The 16S rRNA V4 hypervariable region was targeted by 515F (Parada) 5’-GTGYCAGCMGCCGCGGTAA-3’ (Parada et al., 2016); 806R (Apprill) GGACTACNVGGGTWTCTAAT (Apprill et al., 2015), and the 18S rRNA V9 hypervariable region was targeted by 1391f 5′-GTACACACCGCCCGTC-3′ (Lane, 1991) and EukBr 5′-TGATCCTTCTGCAGGTTCACCTAC-3′ (Medlin et al., 1988). Amplification conditions for 16S V4 were 98 °C for 30 seconds; 30 cycles of 98 °C for 10 seconds, 55 °C for 30 seconds, 72 °C for 30 seconds; and a final extension of 72 °C for 2 minutes. 18S V9 conditions were the same, with the exception of an annealing temperature of 60 °C. Samples were amplified and multiplexed in a 1-step PCR with a dual-indexing scheme (Kozich et al., 2013). Individual samples were run as 50 μL reactions with 2 ng starting DNA, 25 μL NEBNext High-Fidelity PCR Master Mix (New England Biolabs), 0.5 μM forward and reverse primers, prior to pooling and sequencing by the University of Exeter Sequencing Service on the Illumina MiSeq, using v2 chemistry (250 bp paired-end for 16S and 150 bp paired-end for 18S). The sequencing runs included four positive controls (ZymoBIOMICS® Microbial Community DNA standard, lot number ZRC190811) and six negative controls comprising nuclease free water carried through the entire DNA extraction and PCR amplification.

### 2.4. Bioinformatics processing

All bioinformatics and statistical analyses were performed in R v3.6.3. Following sample demultiplexing, reads were quality controlled and processed by the DADA2 pipeline v1.14 (Callahan et al., 2016). Briefly, quality profiles of paired reads were inspected and forward and reverse reads were truncated at 200 bp and 160 bp, respectively, for prokaryotes, and 100 bp for both reads of microeukaryotes. Amplicon sequence variants (ASVs) were then inferred with DADA2’s pooling method to enhance the detection of rare ASVs. Paired reads were merged if they achieved a minimum overlap of 100 bp for prokaryotes and 25 bp for microeukaryotes. To remove off-target sequencing artefacts, final ASVs were only retained for the lengths 250 – 256 bp for prokaryotes and 90 – 150 bp for microeukaryotes. Chimeras were removed and taxonomy assigned to each ASV against the SILVA SSU v138 taxonomic database (Quast et al., 2012). For the microeukaryotic dataset, only ASVs classified by SILVA as eukaryotic were retained and final taxonomic classifications of these ASVs were made by the PR2 v4.12 taxonomic database (Guillou et al., 2012). Accuracy of the taxonomic assignment was assessed in positive controls, with all members of the ZYMO mock community present, as expected (Supplementary Fig. 2).

A phylogenetic tree of ASVs was constructed with Phangorn v2.5.5 (Schliep, 2010) by first generating a neighbour-joining tree, followed by fitting a generalised time reversible substitution model to generate a maximum likelihood tree. The statistical tool Decontam v1.6 (Davis et al., 2018) was used to identify contaminating ASVs by looking at the prevalence in negative controls, with standard parameters baring a 0.5 prevalence threshold. Thus, all sequences found at greater prevalence in negative controls than positive samples were classed as contaminants and were removed from the ASV table.

ASVs and sample data were parsed to Phyloseq v1.30 (McMurdie and Holmes, 2013) for all subsequent quality control and data analyses. To remove sequencing noise, only ASVs that reached a 2% prevalence threshold across samples were retained. Furthermore, any ASVs taxonomically assigned as chloroplasts, mitochondria, eukaryotic or unclassified at kingdom level were removed from the prokaryotic dataset. Additionally, for the microeukaryotic dataset, 14 sequences classified as Craniata were removed, as these most likely represented fish sequences. As a result, the characterisation of fish skin microeukaryotes was limited due to the high levels of contaminating host 18S rRNA sequences (98.6%) in swab samples.

### 2.5. Statistical and data analysis

Alpha diversity metrics were calculated with Phyloseq on counts rarefied to the minimum sequencing depth. The difference between pond sites was statistically tested by Welch’s ANOVA and *post-hoc* pair-wise Games-Howell test, following confirmation of normality. Further testing between sample types utilised lmerTest v3.1-3 (Kuznetsova et al., 2017) to perform a linear mixed-effects model that accounted for pond site as a random effect. A Pearson’s correlation coefficient was used to test for correlation of Chao1 richness and Shannon diversity between sample types.

Beta diversity analysis was performed with compositional data analysis principles. These comprise log-based transformations, which cannot be performed on zero values. Therefore, ASV counts were subjected to a count zero multiplicative replacement method in zCompositions v1.3.4 (Palarea-Albaladejo and Martín-Fernández, 2015). A centred log-ratio (CLR) transformation was then applied to ASV counts with the CoDaSeq package v0.99.6 (https://github.com/ggloor/CoDaSeq). Euclidean distance was calculated on log-ratios and ordinated by PCoA biplot with FactoExtra v1.0.7 (https://github.com/kassambara/factoextra). Statistical differences between pond site and sample type groups were conducted on the Euclidean distance matrix by permutational multivariate analysis of variance (PERMANOVA) and permutation tests for homogeneity of multivariate dispersions, implemented in Vegan v2.5-6 (Dixon, 2003).

Community composition was presented as heat trees of taxon relative abundance with Metacoder v0.3.4 (Foster et al., 2017), utilising a Davidson-Harel layout algorithm. Differential abundance between sample types was assessed by CornCob v0.1 (Martin et al., 2020), utilising the Wald Chi-Squared test and accounting for pond site as a random effect. Core microbiome analysis was performed on ASVs amalgamated to genus level and rarefied to the minimum sequencing depth. Classification of the fish skin core genera was performed with the Microbiome package v2.1 (Lahti and Shetty, 2017) based on a prevalence threshold of 80% and a detection threshold of 0.01% in all swab samples. Heatmaps of core genera and discriminant taxa were depicted as heatmaps of CLR abundance of non-rarefied counts by pheatmap v1.0.12 (https://github.com/raivokolde/pheatmap).

The significance level and false discovery rate of 0.05 was set for all statistical analyses.

### 2.6. Data availability

Raw sequencing reads were deposited in the European Nucleotide Archive under the accession PRJEB46984. Data processing, analysis scripts and final ASV tables are accessible at https://github.com/jamiemcm/Malawi_Tilapia_Microbiomes.

## 3. Results

Following quality control and filtering, the final prokaryotic dataset contained 969,562 reads and 5782 ASVs from all skin swab and pond water filter samples, respectively, collected in this study (67 samples). The eukaryotic dataset comprised 94,611 reads, 1659 ASVs from the 62 samples collected. Full read counts per library, including break down between skin swabs and pond water filters, are available in Supplementary Table 1.

### 3.1. Phytoplankton communities

Compositional approaches (CLR) to beta diversity were applied to explore variation in microbial community composition and abundance of pond water between sites are shown in Figures 1A and 1B. Clear clustering of water samples by pond site was evident, with the position of mean group centroids corresponding to site shown to be significantly different from each other according to PERMANOVA for both prokaryotic (F(6,28) = 34.29, *R*^2^ = 0.88, *p* < 0.001) and microeukaryotic (F(6,28) = 15.12, *R*^2^ = 0.76 *p* < 0.001) communities. Dispersion of the pond water samples collected within each site was relatively small, particularly with respect to prokaryotes (Fig. 1B). However, largely due to pond site 2, dispersion in prokaryotes differed significantly according to permutation tests for homogeneity of multivariate dispersions (Prokaryotes F(6,28) = 3.95, *p* = 0.003; Eukaryotes F(6,28) = 0.87, *p* = 0.53).

**Figure 1:**
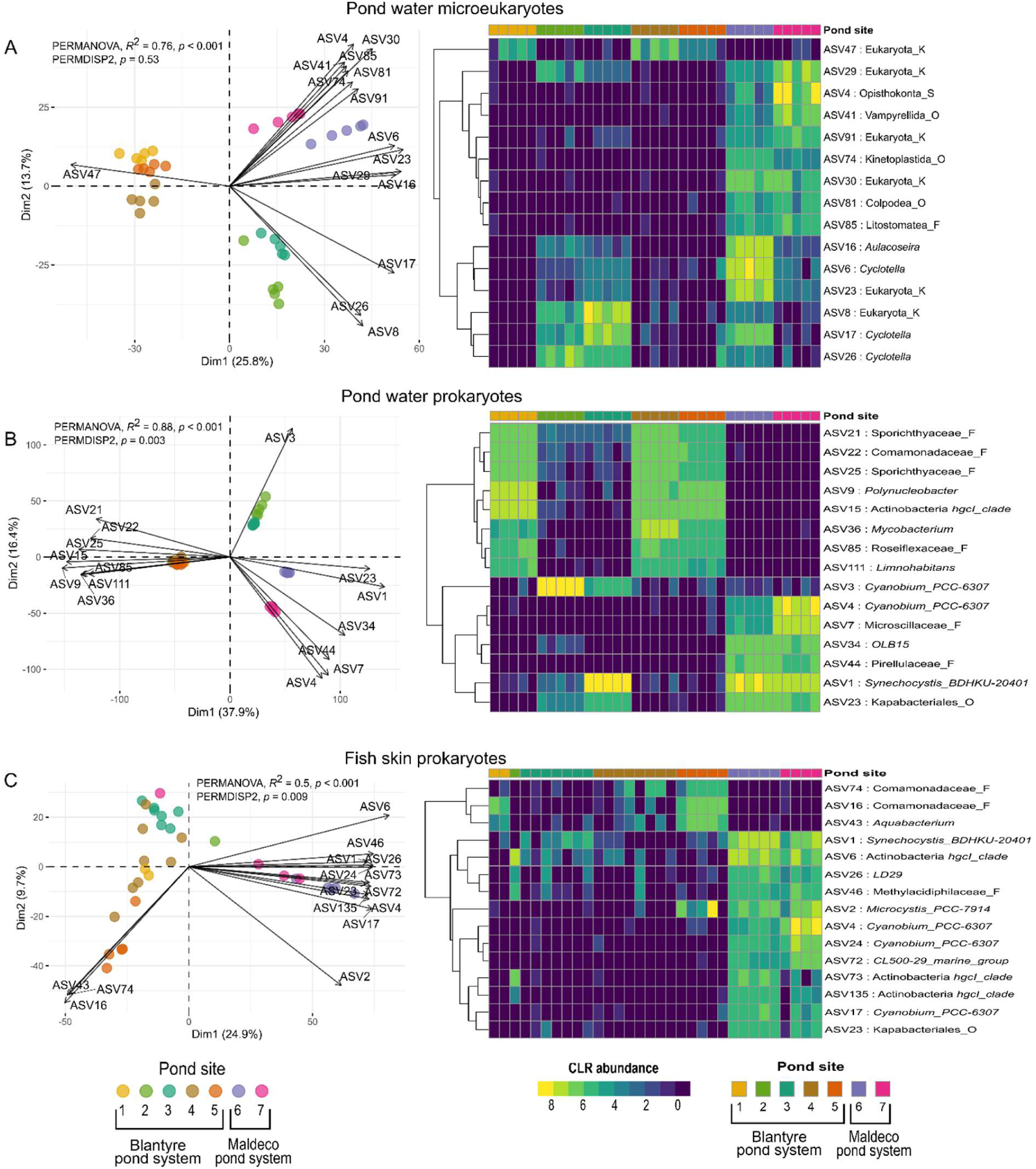
Microbial compositional diversity and abundance of pond water varies significantly by pond site, with trends for inter-site variation consistent across bacterial and eukaryotic communities. Fish skin samples show significant dispersion within pond sites. **Left panel:** Ordination by PCoA biplots on Euclidean distance of log-ratios (Aitchison distance). Points represent samples, coloured by pond site, with arrows for top 15 Amplicon Sequence Variants (ASV) explaining variation between samples. ASV abundances increase in the direction of arrows and arrow length represents magnitude of change. Angles between arrows denote correlation between ASVs (approximately 0° = correlated, < 90° positive correlation, > 90° negative correlation, 90° no correlation). **Right panel:** The centred log-ratio (CLR) abundances of top 15 discriminant ASVs are plotted as accompanying heatmaps, with ASVs ordered according to a hierarchical clustering dendrogram. Labels include ASV number, lowest available taxonomic classification and rank of this classification e.g. “_F” = Family.

While pond location had a strong influence on the separation of pond water samples, the clustering observed in prokaryotes of the tilapia skin was less distinctive (Fig. 1C). There was a significant difference between the mean centroid position of each pond site by PERMANOVA F(6,25) = 4.19, *R*2 = 0.50, *p* < 0.001) and significant dispersion between fish within the same pond (F(6,25) = 5.32, *p* = 0.009).

Specific taxa were associated with driving community separation between pond sites. Figure 1 shows the top 15 contributing taxa plotted as arrows on each biplot, with their CLR abundance depicted in the accompanying heatmaps. In pond water, microeukaryotes included several diatoms (ASV6: *Cyclotella* and ASV16: *Aulacoseira*), the presence of which separated pond site clusters 2,3 and 6,7 from 1,4,5. ASV47: Eukaryota was the major taxon - found at high abundance - discriminating pond site cluster 1,4,5 from the remaining ponds, and a BLASTn search of this ASV revealed 90% similarity to the microalgae *Cryptomonas*. In the pond water prokaryotic community, photosynthetic Cyanobacteria were particularly prevalent in pond site clusters 2,3 and 6,7, with apparently differing *Cyanobium* ASVs (ASV3, ASV4) in each cluster, and a shared *Synechocystis* (ASV1). Pond site cluster 1,4,5 was distinguished by typical freshwater planktonic Proteobacteria (ASV9: *Polynucleobacter*, ASV111: *Limnohabitans* and ASV22: Comamonadaceae), among others. For fish skin prokaryotes, three out of the top 15 discriminant taxa (ASV43: *Aquabacterium*, and ASV16, ASV74: Comamonadaceae) explained the separation of pond cluster 1,4,5 only. Many of these identified taxa shared taxonomic affiliation to the aforementioned prokaryotes of pond water, but were represented by separate ASVs than those previously identified, such as ASV17, ASV24: *Cyanobium*, ASV16, ASV74: Comamonadaceae and ASV6, ASV73, ASV135: Actinobacteria *hgcl clade* (Warnecke et al., 2004).

Alpha diversity metrics gave an insight into species diversity of the pond water samples from different pond sites as determined through assessing community richness (Chao1) and evenness (Shannon diversity and Inverse Simpson diversity) (Fig. S3A,B). Applying Welch’s ANOVA showed a significant difference between both prokaryotic and microeukaryotic communities of each pond site for all diversity metrics (Tab. S2). No correlation was found between prokaryotic and microeukaryotic communities for the mean richness/diversity metric of each pond site (Fig. S3D) (Pearson’s correlation: Chao1 richness *R* = 0.56, *p* = 0.19; Shannon diversity *R* = 0.35, *p* = 0.44; InvSimpson *R* = -0.024, *p* = 0.96).

### 3.2. Microbial niche separation

We used measures of alpha and beta diversity to explore the influence of the environment (pond water) in shaping tilapia skin prokaryotic microbiota. When controlling for pond site as a random effect in linear mixed-effects modelling, ASV richness of the fish skin was found to be significantly lower than pond water (by 503 ± 59.49 ASVs, *R*^2^c = 0.61, *p* < 0.001) (Fig. 2A). Shannon diversity of fish skin and pond water varied according to pond site, however, there was no overall clear separation between the sample types when the aforementioned statistical model was applied (pond water 4.96 ± 0.12, fish skin 4.72 ± 0.15, *R*^2^c = 0.11, *p* = 0.115) (Fig. 2B). Additionally, neither richness nor diversity were correlated between the fish skin and pond water when comparing between pond sites, according to Pearson correlation tests (Chao1 richness *R* = 0.18, *p* = 0.71; Shannon diversity *R* = 0.11, *p* = 0.81) (Fig. 2C,D). Pair-wise comparisons were made of the beta diversity (Aitchison distance) between samples within each pond site (Fig. 2E) and this showed pond water samples clustered closely together, but greater dispersion was apparent between fish skin samples. The largest Aitchison distance values were seen in the comparisons between pond water and fish skin samples, indicating different prokaryotic community structures between these niches. Although these structures were made up of shared taxa, albeit at different abundances, with 4020 of a total 5782 ASVs detected in both pond water and fish skin (Fig. 2F).

**Figure 2:**
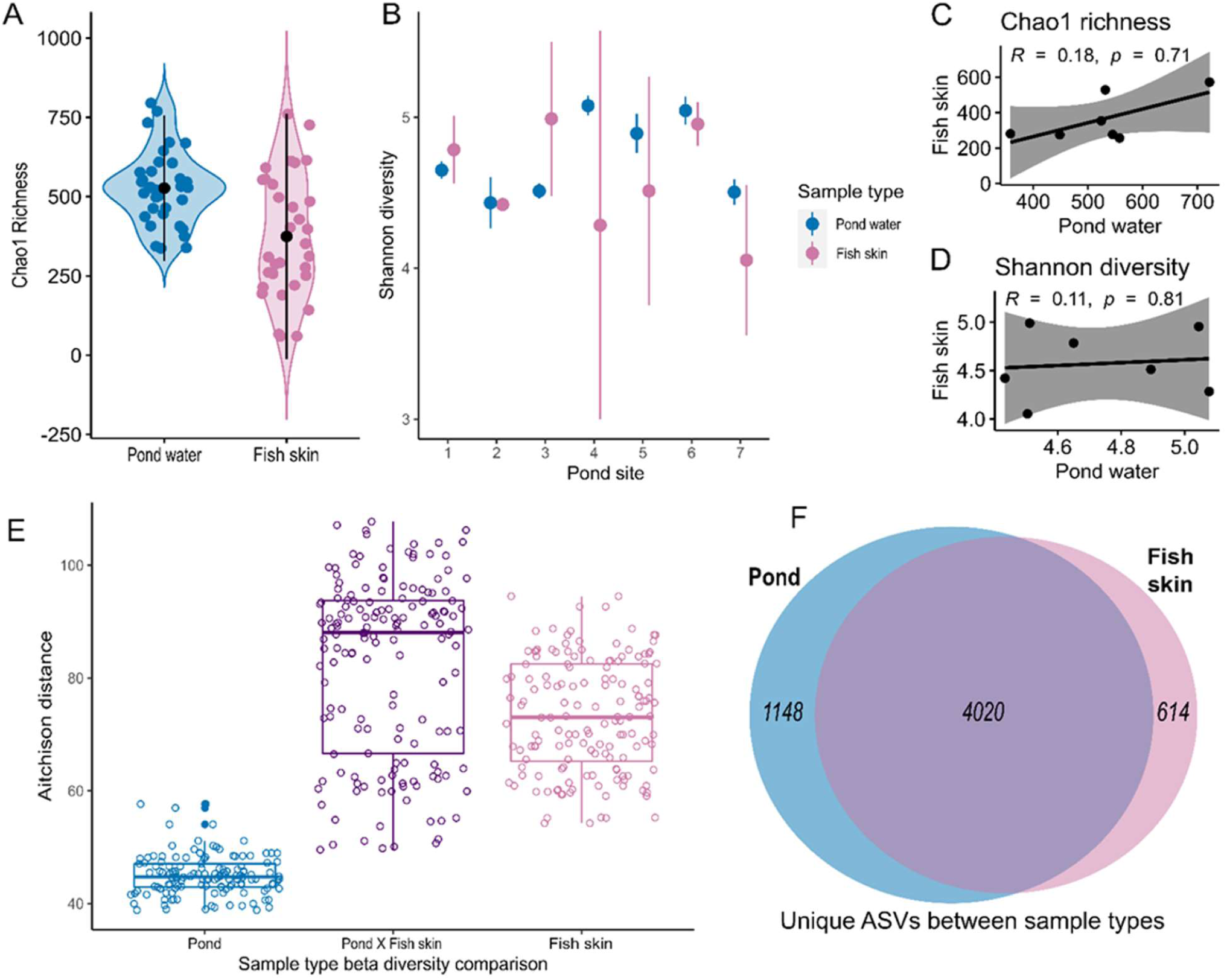
Comparisons of prokaryotic richness and diversity between pond water and fish skin. **A**. Chao1 richness estimates of ASVs per sample, with sample type group means and standard deviations. **B**. Shannon diversity was calculated for each sample and plotted for each pond site as group means and standard deviations. **C, D**. Chao1 richness and Shannon diversity showed no correlation between fish skin and pond water, with points plotted for the mean of each pond site. A regression line of Pearson’s correlation coefficient is plotted, with 95% confidence intervals. **E**. Beta diversity pairwise comparisons of Aitchison distance between samples of pond water, pond water vs fish skin and fish skin, within each pond site. **F**. Number of ASVs unique or shared between pond water and fish skin.

Depicting taxonomic composition of prokaryotic and microeukaryotic communities from skin swab and pond water samples as phylogenetic heat trees (see Fig. S4) illustrates much of the diversity for individual samples is accounted for by rare taxa found at low abundance. The prokaryotic community composition at coarse taxonomic levels was overall very similar between the pond water and skin environments, although divergence emerges at finer taxonomic resolution. For the microeukaryotic community, a far greater overall taxonomic diversity was observed in pond water than on skin, with numerous rare taxa. However, skin diversity was artificially under-sampled due to the over-amplification of tilapia host 18S RNA gene copies.

Taxonomic relative abundance (depicted as dot plots of prokaryotes at class level and microeukaryotes at division level) highlights the differences between pond water and fish skin niches (Fig. 3). According to differential abundance statistical testing, the bacterial classes Gammaproteobacteria and Clostridia were enriched (FDR <0.05) in the fish skin. Pond water by contrast had enriched abundances of Cyanobacteria, Actinobacteria, Bacteroidia, Verrucomicrobiae, Planctomycetes, Kapabacteria and Chloroflexia. Differential abundance testing controlled for pond site as a random effect, however, the degree and consistency of enrichment did vary between pond sites.

**Figure 3:**
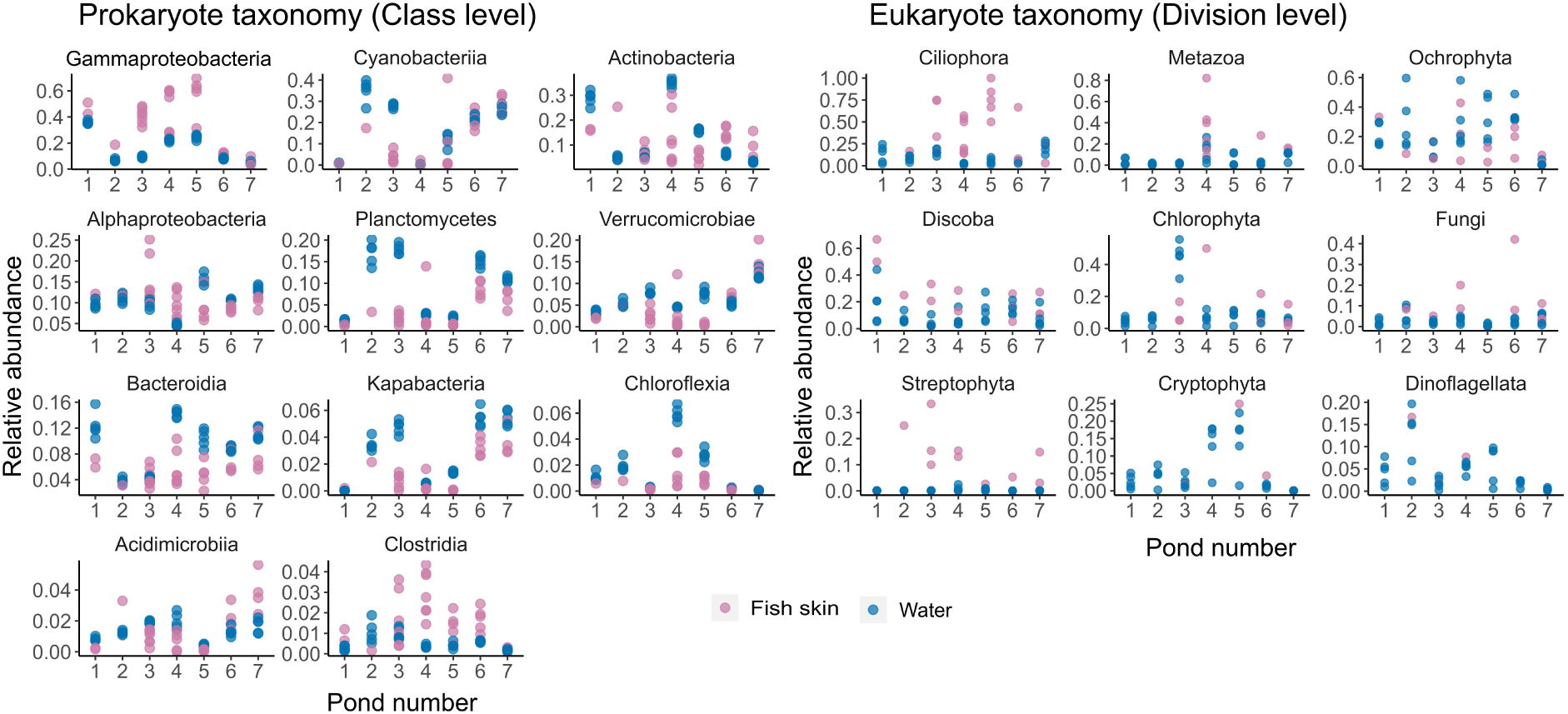
Relative abundance of bacterial (16S) and eukaryotic (18S) taxonomic communities. Each plot reveals the relative abundance for every sample collected at pond sites 1-7, with faceting at the taxonomic levels class (16S) and division (18S). Fish skin and pond water samples are represented in pink and blue respectively. Each facet is scaled independently.

Within these high level taxa, individual prokaryotic taxa were (FDR <0.05) differentially abundant between pond water and the skin (Fig. 4). In general, these taxa followed phylogenetic trends of enrichment, whereby if a taxon was found to be differentially abundant, all other identified taxa within the same phylum were enriched in the same sample type. The pond water was differentially enriched with several taxa associated with key nutrient cycling processes in the aquatic environment, such as the photoautotrophs *Cyanobium, Synechocystis* and *Microcystis*, and the methanotroph *Methylocystis*. Meanwhile, selected ASVs found to be differentially enriched at the skin surface included taxa previously reported as fish microbiome commensals, such as *Cetobacterium*, as well as additional fish related taxa, which in some cases can be associated with diseases, such as *Aeromonas, Pseudomonas, Staphylococcus*, and *Streptococcus*.

**Figure 4:**
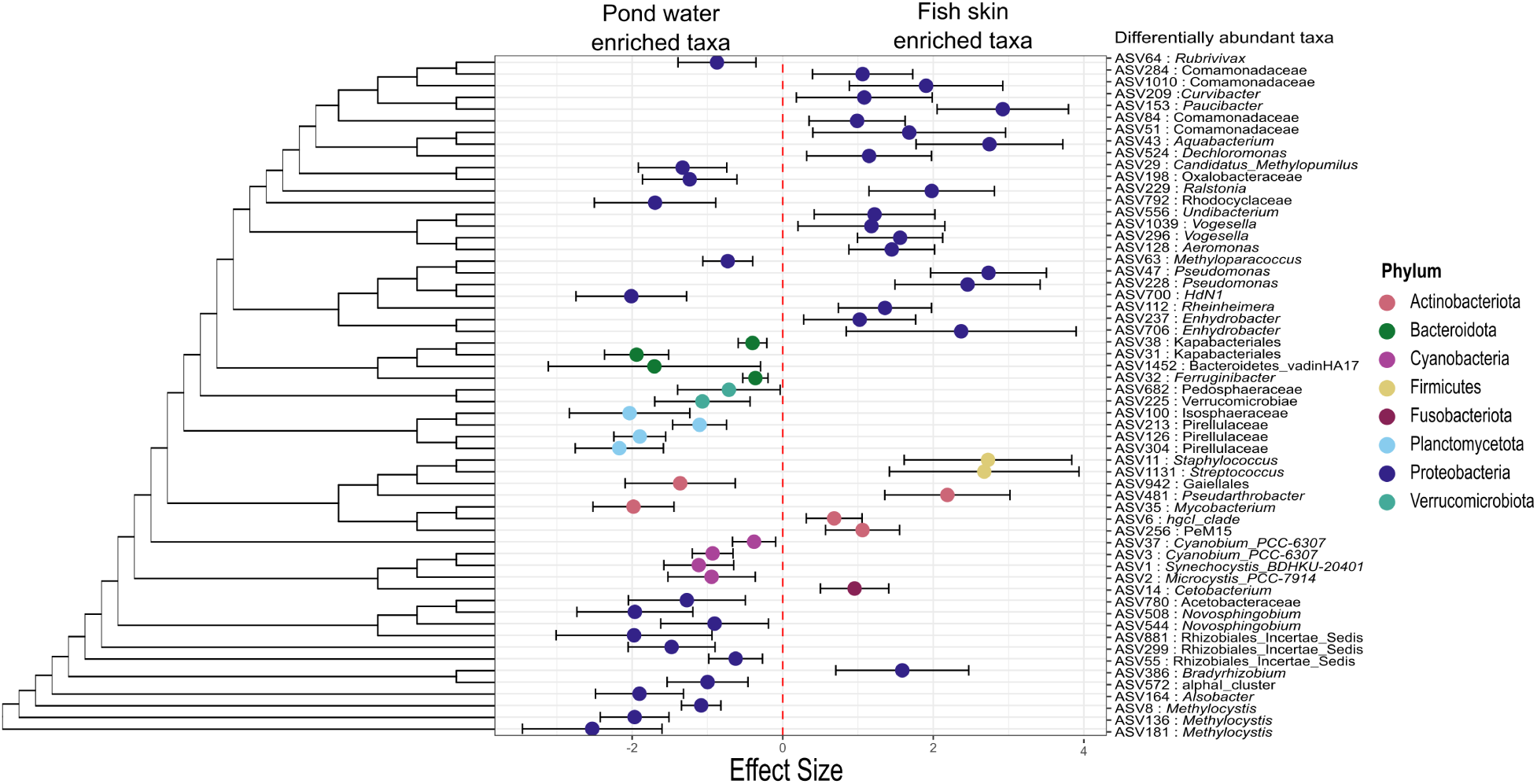
Differentially abundant prokaryotes of pond water and fish skin show phylogenetically conserved trends. The effect sizes and 95% prediction intervals of significant differential abundant taxa (FDR < 0.05) are plotted. Taxa to the left with a negative effect size are enriched in pond water, while taxa with a positive effect size are enriched in the fish skin. Taxa are ordered according to the phylogenetic tree, with labels included for the highest available taxonomic classification of each ASV.

### 3.3. Tilapia species differences

This study featured two tilapia species commonly cultured in Malawi (*Coptodon rendalli* and *Oreochromis shiranus*). No significant difference of prokaryotic community alpha diversity were observed between species (Fig. S5) and while beta diversity did showed potentially unique community structures between species, this only explained 11% of variance. Pond site in contrast explained 50% of variance in beta diversity. Additionally, intra-species dispersion of *C. rendalli* prokaryotic communities (Average distance to median 54.49) was similar to any inter-species dispersion observed between *C. rendalli* and *O. shiranus* at Blantyre (Average distance to median 53.79).

### 3.4. Tilapia skin core microbiome

To further explore specific taxa prevalent within the skin microbial communities we identified 14 prokaryotic core genera of tilapia skin. Abundances of these core genera are depicted for both fish skin and pond water samples in Figure 5. Two of the prokaryotic core genera had a clear enrichment of abundance in the fish skin versus to pond water, namely ASV47: *Pseudomonas* and ASV8731: *Sphingomonas*. The remaining prokaryotic genera were found at high abundance in both pond water and skin samples, despite being classified as part of the tilapia skin core microbiome.

**Figure 5:**
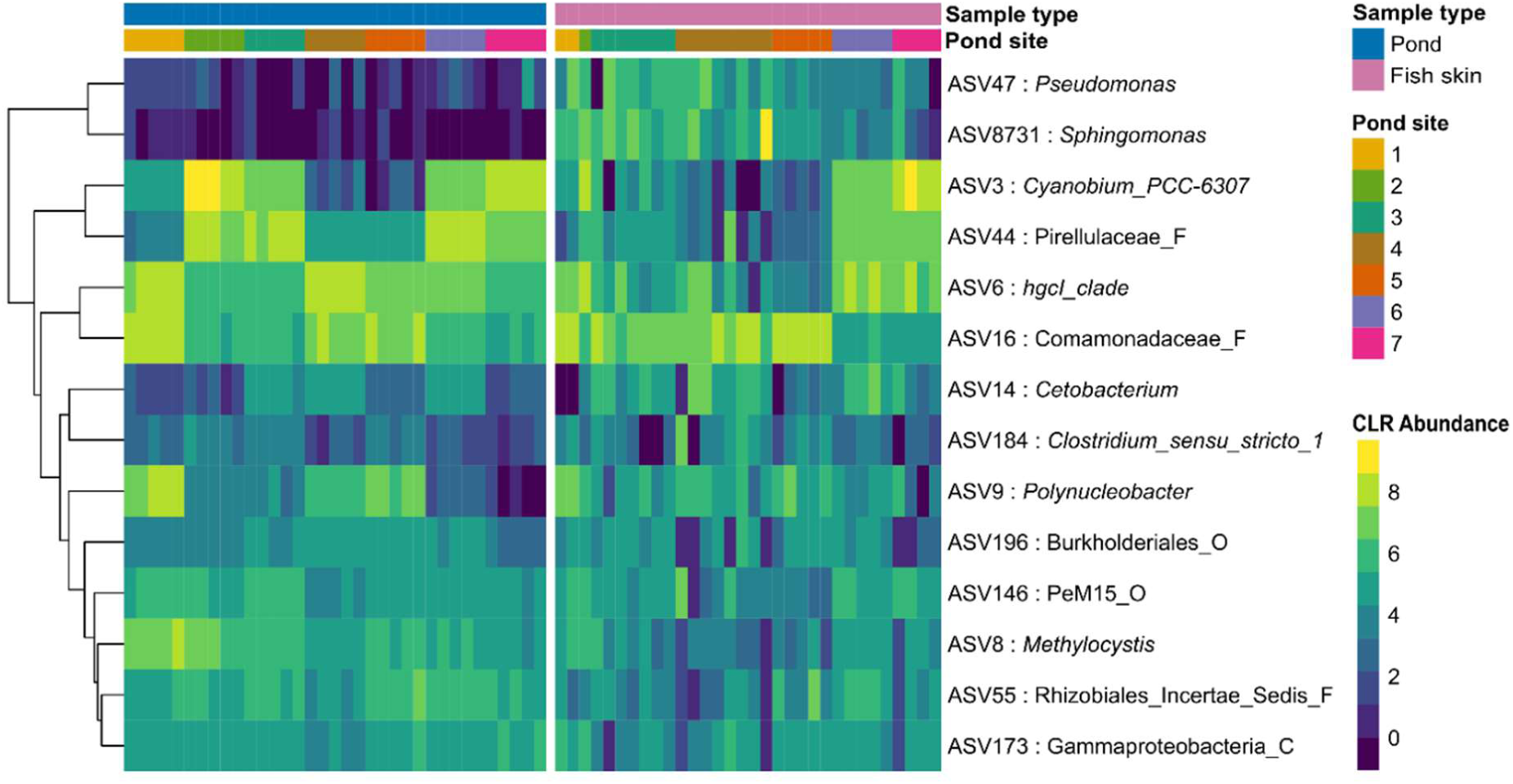
Core bacterial genera of tilapia skin communities. ASVs were amalgamated at genus level, rarefied and core genera were classified across fish samples at an 80% prevalence threshold and 0.01% detection threshold. Abundance counts of core genera were transformed and are presented as non-rarefied compositional log ratios for both pond water and fish skin samples. Abundances were utilised for ordering by a hierarchical clustering dendrogram.

## 4. Discussion

Previous work has highlighted the collective contributions of microbial symbionts, the host and the environment to fish health and disease susceptibility under the pathobiome concept (Bass et al., 2019). Applying this framework to aquaculture production of finfish, the skin mucosal surface microbiome and its direct interface with the environment is likely to play a role in the maintenance of fish health and disease resilience. However, relationships between the microbial assemblages on the skin of fish in culture and their aquatic environment remain poorly established. Here, we characterised the prokaryotic and microeukaryotic communities of the tilapia skin mucosal surface and accompanying water in aquaculture ponds of southern Malawi in the absence of detectable disease to develop a holistic understanding of the relationships between these microbial communities and niches in healthy animals and environments, and against which future studies may assess how microbial dysbiosis contributes to disease onset.

In this study, biogeographic factors played a key role in determining the diversity and structure of pond water microbial communities. Pond location explained 88% of prokaryotic and 76% of microeukaryotic beta diversity separation in microbial abundance profiles. Significant differences in richness and alpha diversity were observed between the seven pond sites. In freshwater ecosystems, both neutral and deterministic processes contribute to the separation of microbial assemblages (Lear et al., 2014; Lee et al., 2013). Interestingly, just over 1% of prokaryotic pond water ASVs were detected in all seven pond sites, suggesting limited species dispersal and/or distinct micro-ecologies between ponds. Numerous environmental selective pressures could play a role in the divergence in ASVs between ponds, such as alternative feeding regimes (Deng et al., 2019), differences in water physiochemistry (Qin et al., 2016) and differences in pond treatments, that can include the use of probiotics (Wu et al., 2016) and manure fertilisers (Minich et al., 2018). Within a pond complex, some of these factors will be conserved, such as weather and water source. Yet microbial community divergence was still observed between ponds in the Blantyre pond complex, with two notable clusters of pond sites (1,4,5 and 2,3).

The tight clustering of pond water samples was concurrent between both prokaryotic and microeukaryotic communities suggesting cross-domain relationships shaped by ecological or environmental processes. This connection has recently been observed in shrimp culture ecosystems, with the deterministic process of homogenous selection largely responsible (Zhou et al., 2021). In this theory, each pond site cluster represents a comparable set of environmental conditions (be it nitrogen, phosphorous or oxygen availability) that exerts strong selective pressures on both prokaryotic and microeukaryotic communities (Zhou and Ning, 2017). Additionally, direct cross-domain ecological interactions may contribute to the observed trends. For instance, phagotrophic protists and their prokaryotic prey have negative interactions (Sherr and Sherr, 2002), while microalgae and bacteria can show all manner of symbiotic relationships, including extensive cross-feeding (Fuentes et al., 2016; Ramanan et al., 2016).

The close proximities of microbial communities of pond water with those in the fish outer mucosal surfaces mean they are physically closely interconnected, yet pond and skin microbiomes clearly differ. Our results demonstrate these differences in prokaryotic community structure, with ASV richness differing significantly, and separation by beta diversity. However, there was no significant difference in alpha diversity, a finding previously reported in freshwater and marine environments (Chiarello et al., 2015; Reinhart et al., 2019; Webster et al., 2018). At finer taxonomic scales further separation between the skin and pond water profiles was seen, and conserved across all ponds sites, with 25 ASVs differentially enriched at the fish skin mucosal surface. The abundances assessed at coarse taxonomic classifications reflected previous reports, namely that Proteobacteia (and in particular Gammaproteobacteria) dominated the fish skin mucosal surface, as seen in a variety of freshwater cichlids (Krotman et al., 2020); reviewed in depth by Gomez and Primm (2021). The next most abundant bacterial classes in the fish skin were Verrucomicrobiae, Bacteroidia and Clostridia. The pond water was similarly dominated by Proteobacteria, followed by Cyanobacteria and Planctomycetes, which is in accordance with a previous report of the bacterioplankton community in Nile tilapia aquaculture ponds in China (Fan et al., 2016).

Despite divergent abundance profiles, there was a high number of taxa shared between pond water and skin mucosa. Only 8% of the total fish skin ASVs were unique to the skin, which contrasts with that for reports on some other fish species. For example, in freshwater river-dwelling mature northern pike, *Esox lucius* L., 36% of skin taxa were not detected in samples of the surrounding water (Reinhart et al., 2019). In a study on freshwater Atlantic salmon, *Salmo salar* L., this figure was 73% (Webster et al., 2018), where fry (8-9 months post-hatch) were sampled from both wild rivers and hatcheries. Both of these studies were in natural aquatic environments and flow-through systems with high water exchange rates which is very different from typical carp and tilapia earthen aquaculture ponds, where daily water exchange rates tend to be very limited, typically a maximum of 20% total pond volume (Nhan et al., 2008). In fact, often in Africa and Asia during dry seasons, due to the lack of water availability, there is no daily water exchange at all in tilapia earthen aquaculture ponds. Such static conditions and high stocking densities may be reflected in a greater microbial crossover between fish skin and pond water. Given the common taxa seen between the tilapia skin and pond water environments, it is noteworthy that we found no correlation in ASV richness or Shannon diversity between the pond and skin niches within each pond site. This finding supports the hypothesis that the skin and pond water niches support uniquely structured microbial communities.

The core microbiome refers to taxa found in the majority of samples which, by inference, may therefore play an important functional role in the microbiome. Fourteen prokaryotic core genera (from a total 770 genera) were identified in tilapia skin, consistent with previously published findings from other studies of fewer than 20 core OTUs on fish skin (reviewed by Gomex and Primm, 2021; Rosado, Pérez-Losada, et al., 2019a). Among the core genera found in the tilapia skin, *Cetobacterium* has been widely reported as a core genus in the gut of freshwater fish (Liu et al., 2016; Sharpton et al., 2021), including tilapia (Bereded et al., 2020; Elsaied et al., 2019). This genus may represent an important functional symbiont, and is reputed to synthesise vitamin B12 and antimicrobial metabolites (Tsuchiya et al., 2007). Other core genera and differentially enriched taxa of the skin are listed in Supplementary Table 3. Ten of the fish skin core genera were also detected at relatively high abundance in pond water. These include *Cyanobium* and *Methylocystis* (two of the most abundant and differentially enriched phytoplankton in pond water), which may have resulted from swab sampling incorporating some residual pond water. Some studies propose only retaining ASVs unique to swab samples and those statistically enriched from water samples to avoid this possible complication (Krotman et al., 2020). This approach, however, risks underestimating diversity and missing key taxa of the fish skin that through mucosal sloughing may still be detected at high abundances in water. The majority of studies make no corrections; instead, acknowledging crossover is inevitable and representative of these niches.

While the current study included two different species of tilapia, geographic location (and the associated environmental factors) of each pond site appeared to be a stronger influence of prokaryotic fish skin communities than any species differences observed between *C. rendalli* and *O. shiranus*. This suggests species is a complicating factor in our study but is of lesser importance when considering the broader trends of microbial community separation between pond water and fish skin. The importance of habitat over host taxonomy has previously been demonstrated for a large-scale study of marine fish gut microbiomes (Kim et al., 2021).

Several of the bacterial genera we found to be differentially abundant in fish skin contained species pathogenic to tilapia, including *Aeromonas* (*hydrophila*) (Dong et al., 2017), *Streptococcus* (*agalactiae*) (Zhang, 2021) and *Pseudomonas* (*fluorescens*) (Hal and El-Barbary, 2020). In addition to these fish skin enriched genera, further potentially pathogenic taxa were detected in pond water and fish skin. Namely, *Plesiomonas* (*shigelloides*) (Liu et al., 2015), *Flavobacterium* (*columnare*) (Dong et al., 2015) and *Acinetobacter* spp. an emerging group of freshwater fish pathogens (Malick et al., 2020). Likewise, among the detected microeukaryotic genera, there were species pathogenic to tilapia, including the parasitic ciliate *Ichthyophthirius multifiliis* (El-Dien and Abdel-Gaber, 2009) and two skin-targeting pathogenic oomycetes, *Aphanomyces invadans* (OIE, 2013) and *Saprolegnia parasitica* (Ellison et al., 2018) but the metabarcoding of short hypervariable regions of marker genes does not allow us to accurately assign species or strain level classifications to determine their pathogenicity. The detected genera also contain numerous non-pathogenic species. The above ‘pathogens’ were all at very low (typically less than 1%) relative abundances in fish skin, and indeed none of these ponds had any reported incidence of disease. This work does not preclude the fact that other pathogens may be present below the limits of detection thresholds or taxonomic resolution. Presence may raise the risk of opportunistic disease as primary or secondary pathogens if environmental stressors create a state of dysbiosis in the fish skin to favour pathobiont propagation, leading to disease onset (Bass et al., 2019).

Contrary to pathogenesis, many of the detected fish skin microbes will exhibit commensal or mutualist relationships with their fish host. For instance, symbiotic bacteria can provide colonisation resistance against pathogens through competition for nutrients and adhesion sites (Legrand et al., 2018). In the microeukaryotic kingdom, ciliates were among the most widely detected taxa of tilapia skin and may offer beneficial roles to the fish by predating upon other microorganisms (Pinheiro and Bols, 2013). Although, the precise functional roles played by symbiotic bacteria and protists of the fish skin remain almost entirely unresolved.

Fish skin microbiomes are inherently variable between populations (Webster et al., 2018), species (Chiarello et al., 2018), individuals in the same environment, and even across different areas of skin (anal, caudal, dorsal and pectoral fins) of the same individual (Chiarello et al., 2015). We observed separation of fish skin communities according to environment (pond site), however inter-individual dispersion within pond sites was considerable, and the degree of dispersion between pond sites was significantly different. This compares to pond water microbiomes showing strong similarities within sites, suggesting the fish skin microbiome is less subjected to environmental influences. This may be due to host factors enabling greater buffering tolerance against environmental directed microbial community assembly. Host genetics is further known to contribute to the inter-individual variation of fish skin communities (Boutin et al., 2014). Additionally, fish age has been seen to influence individual taxa abundances but offers a limited explanation of inter-individual variation at the microbial community level (Rosado et al., 2021). To account for the observed inter-individual variation of fish skin microbiomes we recommend increased fish numbers (6 or more) per treatment/location during sampling campaigns.

In contrast to the variability of fish skin, pond water communities were more consistent across sample sites. At most pond sites, one photoautotroph (*Cyanobium* or *Synechocystis*) was dominant, at up to 20% relative abundance. While *Synechocystis* is well studied as a model organism, little is known of *Cyanobium* and its large contribution to primary production despite being among the most abundant taxa in carp aquaculture ponds (Marmen et al., 2021) and freshwater lakes (Rogers et al., 2021). Additionally, the harmful algal bloom agent *Microcystis* was detected at very high abundance in pond sites 5, 6 and 7, which is concurrent with observations of rich blue-green algae during sampling. *Microcystis* (see Marmen et al., 2021, 2016; Zimba and Grimm, 2003), and its toxin microcystin, are frequently detected in aquaculture ponds and can have toxic effects in tilapia (Abdel-Latif and Khashaba, 2017). Conversely, eukaryotic microalgae, in particular diatoms, contribute positively to the freshwater ecosystem as key primary producers and stabilisers of water quality (Guedes and Malcata, 2012; Li et al., 2017). The barbed spines of some diatoms (*Chaetoceros* spp.), however, can cause gill haemorrhage in saltwater aquaculture (Yang and Albright, 1992). Pond sites 2, 3 and 6 were dominated by several diatoms, including *Cyclotella, Nitzschia* and *Aulacoseira*. In other pond sites, many high abundance ASVs remained unclassified beyond kingdom level, however, BLASTn searches suggested several of these were photosynthetic microalgae and likely contribute to oxygen cycling.

## 5. Conclusions

This study highlights the diversity, structure and variance of the microbial communities found in tilapia skin and pond water, and characterises the microbiomes for ‘healthy’ earthen aquaculture ponds in Malawi. Future studies seeking to establish relationships between dysbiosis and disease states need to take into account the inter-individual variation between fish, and community variance across pond sites that also occurs within the same pond complex. We found a large degree of taxa crossover between fish skin and pond water, some of which may be reflective of swab sampling bias, but also unique microbial communities supported by each niche. The identified core genera and differentially enriched taxa may represent conserved markers of tilapia skin, whose presence and abundance should be considered in future dysbiosis events, albeit in most cases the functional host relation of these taxa at the level of fish skin remains to be determined. Developing a deeper understanding on the microbial communities, particularly those that interface between the aquatic environment and culture species from different geographies, is critical for understanding health risks in aquaculture species as production expands and intensifies, bringing with it an increased risk of dysbiosis and incidence of disease.

## 6. Author contribution statement

Conceptualisation CRT, DB, BT; Fieldwork CRT, DB, CG, JN; Formal analysis JM, DLC; Funding acquisition CRT and DB; Methodology SA, DLC; Supervision CRT, DLC, JDD, BT, JC; Visualisation JM, DLC; Writing JM & CRT; Editing JM, SA, DLC, DB, CG, JDD, CVM, JC, BT and CRT.

## 8. Acknowledgements

We thank, for their contributions to this work, Karen Moore, Audrey Farbos and Paul O’Neill from the Exeter Sequencing Service; Nicola Rogers for project management (Newton Fund project); and John Dowdle for lab management during the practical work.

