## Supplementary materials for "Relationships between pond water and tilapia skin microbiomes in aquaculture ponds in Malawi"

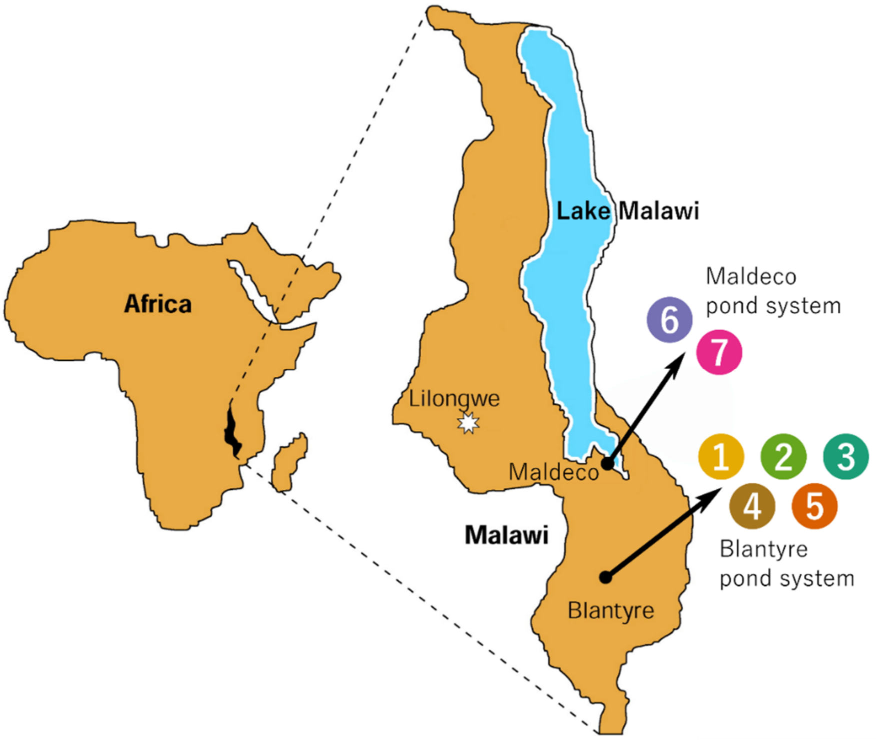


**Supplementary Figure 1: Location of earthen aquaculture pond sites sampled from two pond systems in Malawi.** Map reprinted from The Lancet, Vol. 348, Cetron *et al*., *Schistosomiasis in Lake Malawi*, Pages No. 1274-1278, Copyright (1996), with permission from Elsevier under licence number 5198371305028.

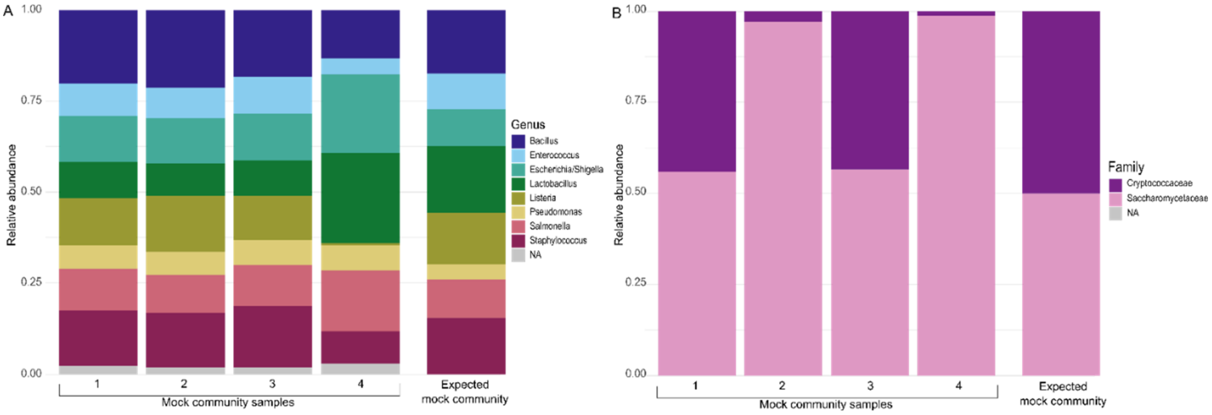


**Supplementary Figure 2: Relative abundance of mock community standards.** A) 16S, B) 18S. Samples 1, 2, 3 represent the ZymoBIOMICS Microbial Community DNA Standard (D6305) which were PCR amplified and sequenced. Sample 4 (ZymoBIOMICS Microbial Community Standard D6300) was carried through the complete sample processing pipeline, including DNA extraction.

**Supplementary Table 1: Sequencing library sizes for 16S and 18S datasets (following removal of non-target sequences such as chloroplasts and fish ribosomal rRNA copies).**

| Measure | Skin swab, 16S Prokaryotes | Pond water filter, 16S Prokaryotes | Skin swab, 18S Microeukaryotes | Pond water filter, 18S Microeukaryotes |
|---|---|---|---|---|
| Total dataset reads | 309,806 | 659,756 | 479 | 294,132 |
| Total dataset ASVs | 4634 | 5168 | 102 | 1650 |
| Median reads per library | 7852 | 18,685 | 12 | 7491 |
| Minimum reads per library | 1490 | 10,550 | 2 | 1322 |
| Maximum reads per library | 25,424 | 34,497 | 58 | 16,180 |

**Supplementary table 2: Hypothesis testing of difference between pond sites by richness and alpha diversity metrics for prokaryotic 16S and microeukaryotic 18S communities. Statistical testing by Welch's ANOVA.**

| Dataset | Metric | ***F* value** | **$\varepsilon^2$ (90% CI)** | ***P* value** |
|---|---|---|---|---|
| 16S | Chao1 | F(6,12) = 9.99 | 0.75 (0.36, 0.84) | < **0.001** |
| 16S | Shannon | F(6,12) = 68.66 | 0.96 (0.90, 0.98) | < **0.001** |
| 16S | Inverse Simpson | F(6,12) = 186.76 | 0.98 (0.96, 0.99) | < **0.001** |
| 18S | Chao1 | F(6,12) = 9.03 | 0.72 (0.31, 0.83) | < **0.001** |
| 18S | Shannon | F(6,12) = 8.39 | 0.71 (0.26, 0.82) | < **0.001** |
| 18S | Inverse Simpson | F(6,12) = 16.07 | 0.83 (0.58, 0.90) | < **0.001** |

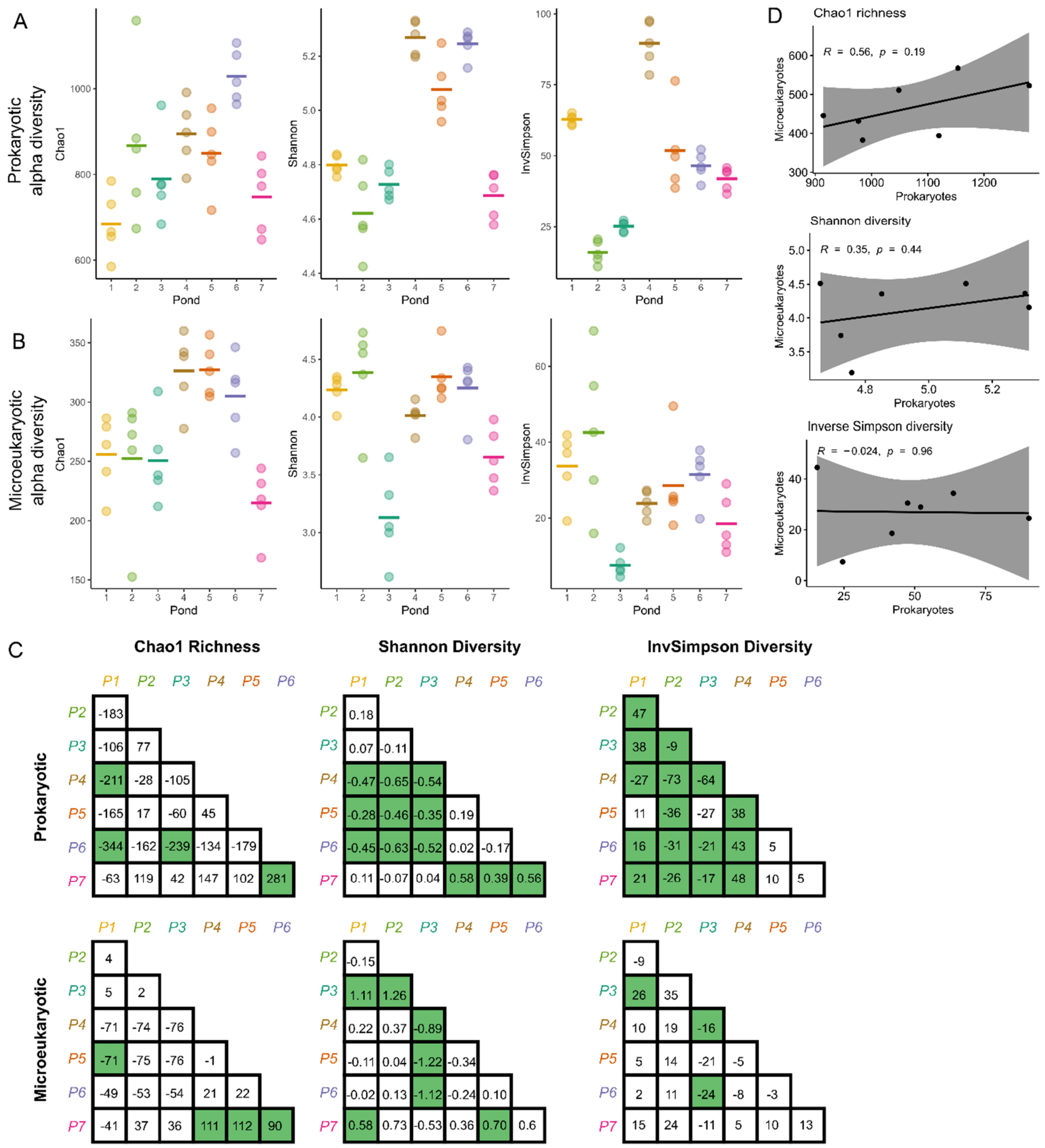


**Supplementary figure 3: Plugin richness and alpha diversity metrics of pond water samples assessing differences between pond sites. A)** Prokaryotic 16S communities, **B)** Microeukaryotic 18S communities. Plugin estimates of richness and alpha diversity by Chao1, Shannon and Inverse Simpson metrics. Group means are displayed as lines. For each metric, significant difference between pond sites was tested by Welch's ANOVA, with results details in Supplementary Table 2. **C)** Pairwise comparisons of diversity metrics, showing mean difference between pond sites with shading indicating a significant difference ($P < 0.05$) according to Games-Howell post-hoc testing. **D)** No correlation in Chao1, Shannon or Inverse Simpson metrics was found between prokaryotic and microeukaryotes communities, with points plotted for the mean of each pond site. A regression line of Pearson's correlation coefficient is plotted, with 95% confidence intervals. All ASV counts were rarefied to equal sampling depth.

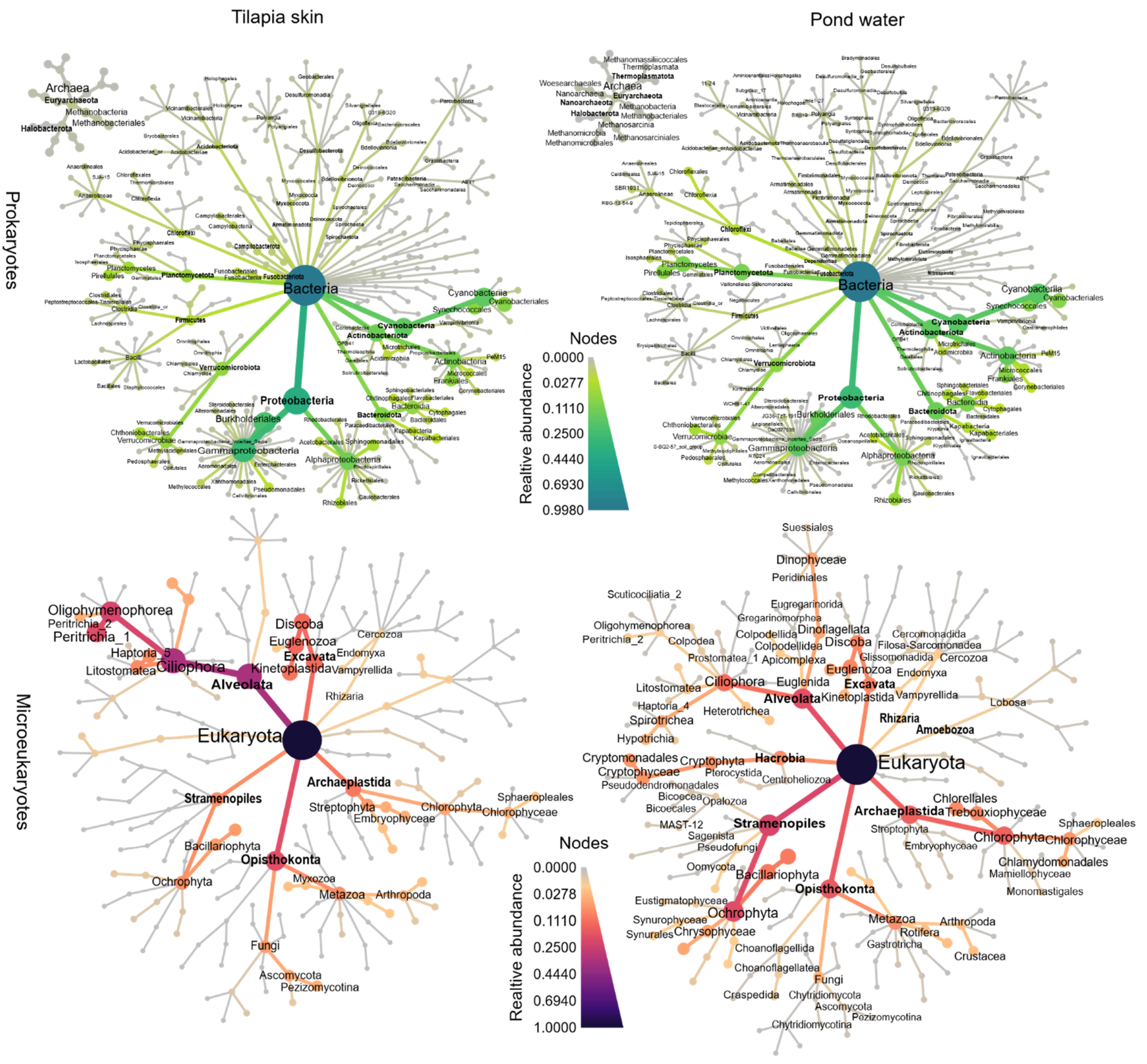


**Supplementary Figure 4: Heat trees depicting taxonomic composition of prokaryotes and microeukaryotes in skin and pond water of Malawian tilapia aquaculture ponds.** Individual taxa present across all samples are represented as nodes up to *Order* taxonomic level. Node size and colour corresponds to the relative abundance of each taxon in sample type groups. Taxonomic labels are included for taxa present in at least 50% of samples from respective sample type groups, apart from eukaryotic skin swabs which have a 10% threshold for taxon labels.

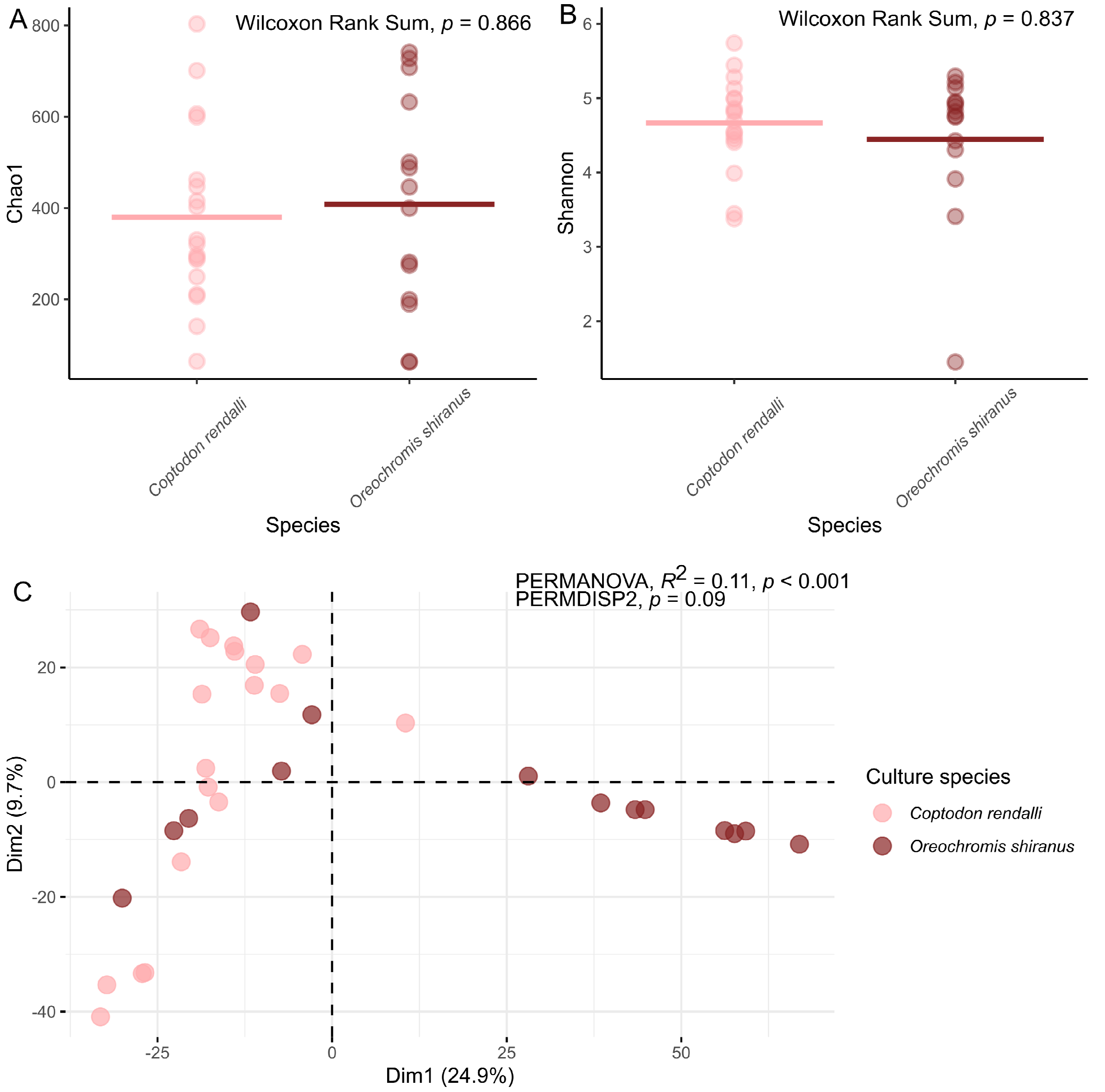


**Supplementary Figure 5: Differences in alpha and beta diversity of skin prokaryotic communities between tilapia culture species *(Coptodon rendalli* and *Oreochromis shiranus)*. A,B)** Rarefied ASV counts of skin swabs for Chao1 richness (A) and Shannon diversity (B), with group means displayed as lines. Statistical difference between sites determined by Wilcoxon rank sum. **C)** Non-rarefied ASVs were subjected to count zero multiplicative replacement prior to compositional transformation by centred log-ratios (CLR) and ordination by PCoA biplots on Euclidean distance of log-ratios (Aitchison distance). Points represent skin swab samples, coloured by species. Difference between pond sites was tested statistically by PERMANOVA, and dispersion within pond sites was determined by permutation tests for homogeneity of multivariate dispersions.

**Supplementary Table 3: Tilapia skin associated prokaryotic taxa with core microbiome and/or differentially abundant status.** Monitoring changes to abundance and variance of these taxa in the fish skin may be useful for future studies to signal dysbiotic events associated with disease.

| ASV | Taxonomy | Reputed function | Previous reports as dominant taxa | Citation |
|---|---|---|---|---|
| 6 | (Actinobacteria) *hgcI clade* | Unresolved, common bacterioplankton | Skin; *Mesonauta festivus, Mylossoma duriventre, Serrasalmus rhombeus* | (Sylvain et al., 2020) |
| 9 | *Polynucleobacter* | Unresolved, common bacterioplankton | Skin; *Esox lucius, Danio rerio* | (Hu et al., 2021; Reinhart et al., 2019; Sangwan et al., 2016) |
| 11 | *Staphylococcus* | Pathogenic and non-pathogenic commensals | Gut; *Lateolabrax maculatus*. Skin and gut; *Hypomesus nipponensis* | (Duan et al., 2021; Rosenstein and Götz, 2012; Soriano et al., 2018) |
| 14 | *Cetobacterium* | Synthesis of vitamin B-12 and antimicrobial metabolites | Gut; *Oreochromis niloticus, Arapaima gigas*. Skin; *Acanthobrama lissneri, Capoeta damascina, Carasobarbus canis, Garra jordanica, Oxynoemacheilus insignis, Coptodon zillii, Oreochromis aureus, Sarotherodon galilaeus, Danio rerio* | (Hu et al., 2021; Krotman et al., 2020; Ramírez et al., 2018; Suphoronski et al., 2019; Tsuchiya et al., 2007) |
| 43 | *Aquabacterium* | Unresolved | Gut; *Salmo salar*. Skin; *Gambusia affinis* | (Gupta et al., 2019; Leonard et al., 2014) |
| 44 | Pirellulaceae | Nitrification | Gut; *Haplotaxodon trifasciatus, Haplotaxodon microlepis, Plecodus straeleni, Perissodus microlepis* | (Baldo et al., 2015; Kellogg et al., 2016) |
| 47, 228 | *Pseudomonas* | Numerous opportunistic fish pathogens (alongside non-pathogenic environmental strains) | Skin; *Salmo salar, Mugil cephalus, Lutjanus campechanus, Cynoscion nebulosus, Cynoscion arenarius, Lagodon rhomboides* | (Larsen et al., 2013; Minniti et al., 2017; Oh et al., 2019) |

| | | | | |
|---|---|---|---|---|
| 51, 84, 284, 1010 | Comamondaceae | Unresolved | Gut; *Seriola lalandi*, *Danio rerio.* Skin and gut; *Hypomesus nipponensis* | (Park and Kim, 2021; Sharpton et al., 2021; Soriano et al., 2018) |
| 112 | *Rheinheimera* | Antimicrobial synthesis | Skin; *Salvelinus fontinalis* | (Boutin et al., 2014; Chen, 2010) |
| 128 | *Aeromonas* | Numerous fish pathogens | Gut; *Salmo salar*, *Ictalurus punctatus*, *Micropterus salmoides*, *Lepomis macrochirus* | (Chen et al., 2019; Larsen et al., 2014; Navarrete et al., 2008) |
| 146, 256 | (Actinobacteria) PeM15 | Unresolved | Gut; *Hypophthalmichthys nobilis* | (Meng et al., 2021) |
| 153 | *Paucibacter* | Degradation of cyanotoxins | Gut; *Danio rerio* | (Rapala et al., 2005; Sharpton et al., 2021) |
| 184 | *Clostridium (sensu stricto 1)* | Typical gut commensals | Gut; *Oreochromis niloticus*, *Arapaima gigas*, *Hypophthalmichthys molitrix*, *Hypophthalmichthys nobilis* | (Li et al., 2017; Lopetuso et al., 2013; Ramírez et al., 2018; Wu et al., 2021) |
| 209 | *Curvibacter* | Iron oxidation | Gut; *Ctenopharyngodon idellus* , *Megalobrama amblycephala*, *Carassius auratus*, *Cyprinus carpio*, *Hypophthalmichthys molitrix*, *Hypophthalmichthys nobilis* | (Gülay et al., 2018; Li et al., 2014) |
| 237, 706 | *Enhydrobacter* | Unresolved | Skin; *Danio rerio*, *Mesonauta festivus*, *Mylossoma duriventre*, *Serrasalmus rhombeus*, *Gambusia affinis* | (Hu et al., 2021; Leonard et al., 2014; Sylvain et al., 2016) |
| 296, 1039 | *Vogesella* | Unresolved | Gut; *Astyanax mexicanus* | (Ornelas-García et al., 2018) |
| 386 | *Bradyrhizobium* | Associated with lipid metabolism | Gut; *Salmo salar*, *Paralichthys olivaceus* | (Dvergedal et al., 2020; Niu et al., 2020) |
| 481 | *Pseudarthrobacter* | Unresolved | Unreported | |
| 524 | *Dechloromonas* | Polyphosphate accumulation | Aquaculture pond sediment | (Petriglieri et al., 2021; Zhang et al., 2020) |

| | | | | |
|---|---|---|---|---|
| 556 | *Undibacterium* | Fatty acid and lipid metabolism | Gut; *Oreochromis niloticus.* Skin; *Colossoma macropomum* | (Kim et al., 2014; Sylvain et al., 2016; Wu et al., 2021) |
| 1131 | *Streptococcus* | Numerous fish pathogens | Skin and gut; *Hypomesus nipponensis* | (Mishra et al., 2018; Soriano et al., 2018) |
| 8731 | *Sphingomonas* | Glycosphingolipid synthesis for immune stimulation | Gut and gill; *Oreochromis niloticus.* Skin; *Gambusia affinis* | (Leonard et al., 2014; Wei et al., 2010; Wu et al., 2021) |
